# Extreme distribution of deleterious variation in a historically small and isolated population - insights from the Greenlandic Inuit

**DOI:** 10.1101/061440

**Authors:** Casper-Emil T. Pedersen, Kirk E. Lohmueller, Niels Grarup, Peter Bjerregaard, Torben Hansen, Hans R. Siegismund, Ida Moltke, Anders Albrechtsen

**Affiliations:** Department of Biology, Section for Computational and RNA Biology, University of Copenhagen, 2200 Copenhagen N, Denmark; Department of Ecology and Evolutionary Biology, University of California, Los Angeles, CA 90095, United States; Department of Human Genetics, David Geffen School of Medicine, University of California, Los Angeles, CA 90095, United States; The Novo Nordisk Foundation Center for Basic Metabolic Research, Section of Metabolic Genetics, Faculty of Health and Medical Sciences, University of Copenhagen, 2100 Copenhagen E, Denmark; National Institute of Public Health, University of Southern Denmark,1353 Copenhagen K, Denmark

## Abstract

The genetic consequences of a severe bottleneck on genetic load in humans are widely disputed. Based on exome sequencing of 18 Greenlandic Inuit we show that the Inuit have undergone a severe ~20,000 yearlong bottleneck. This has led to a markedly more extreme distribution of deleterious alleles than seen for any other human population. Compared to populations with much larger population sizes, we see an overall reduction in the number of variable sites, increased numbers of fixed sites, a lower heterozygosity, and increased mean allele frequency as well as more homozygous deleterious genotypes. This means, that the Inuit population is the perfect population to examine the effect of a bottleneck on genetic load. Compared to the European, Asian and African populations, we do not observe a difference in the overall number of derived alleles. In contrast, using proxies for genetic load we find that selection has acted less efficiently in the Inuit, under a recessive model. This fits with our simulations that predict a similar number of derived alleles but a true higher genetic load for the Inuit regardless of the genetic model. Finally, we find that the Inuit population has a great potential for mapping of disease-causing variants that are rare in large populations. In fact, we show that these alleles are more likely to be common, and thus easy to map, in the Inuit than in the Finnish and Latino populations; populations considered highly valuable for mapping studies due to recent bottleneck events.

## Introduction

Predictions about the consequences of a small population size on genetic variation are among the most fundamental theoretical predictions in population genetics ^1–3^. Small populations are more affected by drift than are large populations; therefore, small populations are predicted to carry lower genetic diversity ^1^. Additionally, natural selection acting against deleterious alleles is predicted to be less efficient in small populations ^2^. Finally, the genetic load, defined as the reduction in fitness caused by deleterious variation ^4^, is predicted to be larger in small populations^1^. With the recent advent of genome-wide sequencing data from humans, it has become possible to test these predictions in human populations and numerous studies have pursued this by comparing the distributions of deleterious alleles across human populations ^5–7^. Most of these studies have focused on genetic consequences of the bottlenecks, i.e. rapid large decreases in population sizes that all non-Africans populations went through during the Out-Of-Africa (OOA) dispersal. These studies all agreed that the European population ^6–9^ and other non-African populations ^5^ carry lower levels of genetic diversity compared to the African populations ^10,11^ and thus have a higher genetic load assuming mutations have a recessive effect. However, the studies ^7,9,10,12^ disagreed in their conclusions about the extent to which genetic load vary across populations assuming mutation have an additive effect and to whether selection is less efficient in small populations.

There is no direct way of calculating genetic load from genetic data from natural populations ^11^. Instead, summary statistics are used and the conclusions based on different statistics in different studies vary. On one hand, a recent study by Do et al. ^8^ found no difference between the non-African and African 1000 Genomes populations using the R_X/Y_ statistic, which is monotonically related to the difference in number of derived alleles between the two populations. In line with this finding, two other studies found no significant difference in the number of derived alleles per genome between individuals of European descent and individuals of African American descent ^7,8^. Based on this observation, Simons et al. ^7^ concluded that Europeans and African Americans carry the same amount of genetic load and supported by simulations, they furthermore concluded that genetic load is not affected much by recent population size changes. On the other hand, other studies found slightly, but significantly, more derived deleterious alleles per individual in the European population than in African populations ^6,9^ implying that the OOA bottleneck has led to a small increase in genetic load in the European population. Similarly, in a recent study of seven populations, Henn et al. ^5^ reported that the non-African populations on average harbor slightly more derived alleles among alleles with large Genomic Evolutionary Rate Profiling (GERP) scores. Furthermore, Henn et al. ^5^ reported a small, but significant, difference in load across the seven populations under an additive effect model when load is estimated using selection coefficients approximated based on GERP scores.

The consequences of smaller population sizes on the efficacy of selection in humans was first investigated in a study of exome data from 15 African Americans and 20 Europeans ^13^. That study showed that the proportion of SNPs that are non-synonymous, and thus likely deleterious, is larger in Europeans than in African Americans. This observation was initially interpreted as being, in part, due to less efficient selection during the OOA bottleneck ^5^ combined with the recent influx of nonsynonymous mutations during the recovery of the bottleneck. Several later studies have come to similar conclusions ^5,9,11^. In contrast, based on their results about genetic load described above, Do et al. ^8^ concluded the efficacy of selection was not reduced due to the reduced population size during the OOA bottleneck ^7,8^. A recent review ^11^ tried to reconcile the different observations and conclusions by pointing out that different studies have focused on different definitions of the efficacy of selection and use different metrics to quantify it. The review ends by calling for descriptions of empirical patterns of deleterious mutations in other human populations than Europeans and Africans with different population size histories to shed further light on the questions about the effect of small size on selection and load. Motivated by this, we here analyze patterns of deleterious mutations in the Greenlandic Inuit (GI) population based on exome sequencing data from 18 GI individuals.

The GI came to Greenland less than 1000 years ago ^14,15^ and a previous study has shown that their ancestors split from the closest large old-world population, the Han Chinese (CHB) population, some ~23K years before present with only a limited amount of subsequent gene flow ^16^. The study also showed that the GI population has a very small effective population size ^16^. This suggests that the GI population has likely been small and isolated for a long period of time after the OOA dispersal took place. Analyses of the GI population could therefore be particularly important for resolving the open questions about whether a small effective population size has had any effect on genetic load and selection in human populations.

In this paper we first show that the GI population is indeed one of the smallest populations in the world and therefore valuable to analyze. Then we compare different proxies for genetic load between the GI and Europeans, as well as investigate several additional measures related to efficacy of selection. And based on our results we discuss the effect of a small population size on load and selection. Finally, we present analyses that investigate the consequence of our findings to disease genetics, which show that the GI population has great potential for leading to discoveries of new disease related genetic variants that have been missed in large scale GWAS in Europeans and other large populations.

## Materials and Methods

### Exome Datasets

We based all of our analyses of the GI population on the high-depth whole exome sequencing data from Moltke et al. ^17^. This dataset consists of data from 9 Greenlandic trios, however we restricted our analyses to data from the 18 parents. The GI data was both analyzed alone and together with other datasets: 1) whole exome sequencing data from 18 unrelated Utah residents with Northern and Western European ancestry (CEU), 18 Han Chinese individuals from Beijing (CHB) and Yoruban individuals from Ibadan, Nigeria (YRI) from the 1000 Genomes project ^18^, 2) an exome dataset which includes Mayans, Mbuti and Cambodians from Henn et al. ^5^ and 3) called genotypes from the 1000 Genome low-coverage data. The analyzed datasets are described in detail below.

*GI dataset* This dataset consists of high-depth exome data from 18 GI individuals. To generate it, Moltke et al. ^19^ performed SNP and indel calling followed by genotype calling for the exome data from the 18 Greenlandic samples using Samtools version 0.1.18 ^20^. Reads with a mapping quality lower than 30 as well as bases with a base quality lower than 20 had been removed. SNPs had been called using standard settings and genotypes were called based on the highest genotype likelihood. We used VCFtools version 0.1.11 ^21^ for filtering. We removed SNVs where all individuals were heterozygous, which are likely genotype errors. In addition, we removed sites with sequencing depth lower than 10 for all individuals or higher than 500. The resulting dataset comprise 133.808 SNVs within the exome. This data set was used solely for identifying absolute numbers of SNVs across a number of functional categories and for inferring population size changes though time.

### Combined exome dataset

This dataset consists of exome data from both the GI and the CEU population. Further, this dataset also includes exome data from 18 Han Chinese individuals in Beijing (CHB) and 18 Yoruban individuals in Ibadan, Nigeria (YRI). To generate it, we performed joint SNP and genotype calling for the exome data from both populations using ANGSD ^22^ under the Samtools ^18^ genotype likelihood model. Reads with a mapping quality lower than 30, as well as bases with a base quality lower than 20, were removed. SNVs were called using a likelihood ratio test ^22^ with a *P*-value cutoff of 10^−6^ and genotypes were called based on the highest genotype likelihood. We required a minimum depth of 10 for calling genotypes and removed sites with missing genotypes and sites, which were triploid when including the ancestral allele. This dataset comprises 295.065 SNVs. SFS comparison of 1000G genotype calls from the low depth data and our genotype calling from the exome data for the same individuals revealed no differences in proportions (Figure S7). The datasets was polarized using the chimp data available within Seattleseq 138 annotation. For each pairwise comparison only the sites that are polymorphic in the two populations were used.

### Dataset from 1000 genomes low-depth whole genome sequencing

This dataset consists of data from five of the 1000 genomes population samples^18^. Finns from Finland (FIN), Peruvians from Lima, Peru (PEL), Gujarati Indians from Houston, Texas (GIH), Utah Residents (CEPH) with Northern and Western Ancestry (CEU), Yoruba in Ibadan, Nigeria (YRI) and Han Chinese in Bejing, China (CHB). To generate the dataset, we used VCF files with genotypes calls from the complete phase 3 1000genomes dataset. From this dataset we extracted 11 unadmixed individuals from each of the 5 populations. Because of apparent admixture in the Peruvians, we reduced the sample size for each population to 11 individuals. Unadmixed individuals from PEL were selected based on inferred admixture proportions (<5%) using ADMIXTURE. The final dataset comprised 401.821 SNVs within the exome.

### Exome sequencing data for Mayans, Mbuti and Cambodians

This dataset consists of data from six samples from each of the populations Mayan, Mbuti and Cambodian. To generate it, we downloaded VCF files with called genotypes for individuals from seven populations recently made publicly available by Henn et al. ^5^. From this dataset we extracted data from 6 Mbuti Pygmies, 6 Mayan Indians and 6 Cambodians. More specifically, following Henn et al.^5^ we excluded two of the 8 Mayan samples due to admixture, leaving only 6 Mayan samples. To ensure comparability between the SFSs inferred from the different populations, we included only 6 randomly chosen individuals from each of the other populations. The ancestral allele in each dataset was obtained separately using the Pantro2 chimp allele. The final dataset contains genotype data for 194.278 SNVs.

### Annotation of variants

We divided variant sites into four categories based on the functional category annotated to the derived allele: synonymous, a combined category of non-coding exons (NCE) and the leading and trailing untranslated region (UTR), missense and loss of function (LoF).

Variant sites which belong to other functional annotations were excluded from further analysis. We assigned a variant to the putatively most deleterious annotation category when there were multiple splice variations. We also included an additional category for loss of function where all isoforms are annotated to be loss of function (denoted LoF^a^), meaning that these sites are more likely to be deleterious than the sites in the LoF category. The five categories thus range from mutations that are expected to be neutrally evolving to mutations that are expected to be highly deleterious.

We also divided variants in categories using GERP scores ^23^. These GERP score measures conservation across a phylogeny of 35 mammalian species excluding humans ^5^. Specifically, GERP scores represent the deficiency in numbers of substitutions in functional loci compared to that of the number of substitutions seen in neutral DNA. This discrepancy is then regarded as a sign of functional constraint and thus lower degrees of substitution saturation across the phylogeny will reflect higher levels of purifying selection ^5,24^. We used the same approach as Henn et al. ^10^: we retrieved the GERP scores from the UCSC browser and grouped variants into four categories according to their GERP score, and thus how deleterious they are predicted to be. The categories are “neutral” (GERP <2), “moderate” (2 ≤ GERP < 4), “large” (4 ≤ GERP < 6) and finally “extreme” (GERP ≥ 6).

### Population size inference

We performed inference of population size over time on the GI dataset using the site frequency spectrum (SFS) based method called stairway plot ^25^. We performed the analyis using default settings.

### Genetic load approximating summary statistics the number of derived alleles

We used two different proxies for genetic load based on counts of derived alleles: the total number of derived alleles per individual, which has previously been used to quantify load under an additive model ^9^ and the number of homozygous-derived genotypes per individual, which can equivalently be used to quantify load under a recessive model.

### The GERP score load

We also used what we will denote GERP score load. To calculate this, we used the GERP-based grouping of deleterious sites described above and translated the three nonneutral GERP score categories into values of selection coefficients as suggested by Henn et al. ^10^. Specifically, the extreme category was assigned *s* = 1×10^−2^, the large category was assigned *s* = 4.5×10^−3^, while the moderate category was assigned *s* = 4.5×10^−4^. These assignments were then used in the equation from Kimura^1^

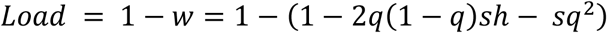

which is based on the underlying model where the fitness of each genotype is determined as 1, 1-*hs* and 1-*s* for AA, Aa and aa, respectively. We used this equation assuming two models each with an extreme level of dominance. In the first model *h* was set to 0.5, i.e. we assumed alleles to act additively. In the other model *h* was set to 0, i.e. we assumed deleterious alleles to be completely recessive. To estimate standard error for our GERP score loads estimates we used a weighted uneven block jackknife, with block sizes of 5Mb to correct for correlation among neighboring sites ^26^.

### π_var_ calculation

We calculated the nucleotide diversity for the variable sites, π_var_, for all the populations included in the SFS comparison in Figure 2. We used the following equation for our calculations:

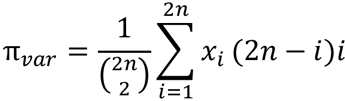

where x_i_ is the proportion of sites in the ith category of the site frequency spectrum and *n* the number of individuals included in the analysis. We calculated π_var_ rather than π for all sites, because it is not possible to calculate π for all sites from the 1000 genomes genotype calls since there is not information for the invariables sites.

### Simulations

We performed simulations using the forward-time simulation software Selection on Linked Mutations (SLIM) ^27^. We used the population size and the demographic history estimated for the GI and the CHB by Fumagalli et al. ^16^. We initiated simulations with a haploid population size of 9395 at 29459 generations before the sampling time (present time), allowing sufficient time for saturation of neutral mutations. During the OOA bottleneck population size was reduced to 5443 haploid genotypes. The CHB population did not change population size up until present time. To simulate the additional bottleneck, which the GI population underwent after splitting from CHB, we introduced an additional reduction in population size 930 generations before the present, resulting in a population size of 1550 haploid individuals. We sampled 36 haploid genotypes from each of the simulated GI and CHB populations, allowing us to capture comparable levels of genetic diversity as the real data where we have 18 diploid samples from each. For this simulated demographic scenario, we chose to simulate many sites under selection: 90% were simulated as neutral mutations and 10% as being under selection. For the deleterious mutations, we varied the dominance coefficient (*h*) and the selection coefficient (s) to mimic the effects of three modes of inheritance: additive (*h*=0.5), near-recessive (*h*=0.1) and recessive (*h*=0). Furthermore, for each of these modes, we varied the selection coefficient, *s*, across four levels, reflecting increasingly harmful functional effects (*s*=0.0002; *s*=0.002; *s*=0.02; *s*=0.2). Note, that because SLIM have a different underlying model of selection implemented than the one we mentioned above, hence the selection coefficients provided here.

Each of the scenarios, i.e. combination of *h* and *s* value, was simulated 10000 times on a sequence of 100k sites using a recombination rate of 1.2 × 10^−8^ bp. and mutation rates of 1.38 × 10^−8^ per generation. Both numbers of derived alleles and numbers of homozygous-derived genotypes are reported excluding mutations that were fixed in both populations. Furthermore, we simulated a scenario where all mutations were set to neutral in GI after the split between GI and CHB. For this simulation, we sampled 36 haploid genotypes both in the pooled population before the split and in each of the GI and CHB populations.

### Allele frequencies in GI compared to Europeans, East Asians, Finns and Latinos

To compare allele frequencies for shared alleles across populations, we used the information available for more than 9.3 million sites based on exome data in the ExAC browser. This dataset has sufficient number of samples for European and East Asian populations to allow reliable calculation of rare allele frequencies.

## Results

### Inference of demography and SFS

First we inferred population size of the GI population over time from exome data for 18 GI using the stairway plot method ^25^. The results suggest that the GI population experienced a marked decrease in size some 22-24 Kya (Figure 1), remained small for more than 20,000 year and only recently started to increase in size. We note that there is a considerable amount of uncertainty for recent population size estimates, which makes it difficult to assess the extent of the recent increase.

**Figure 1.**
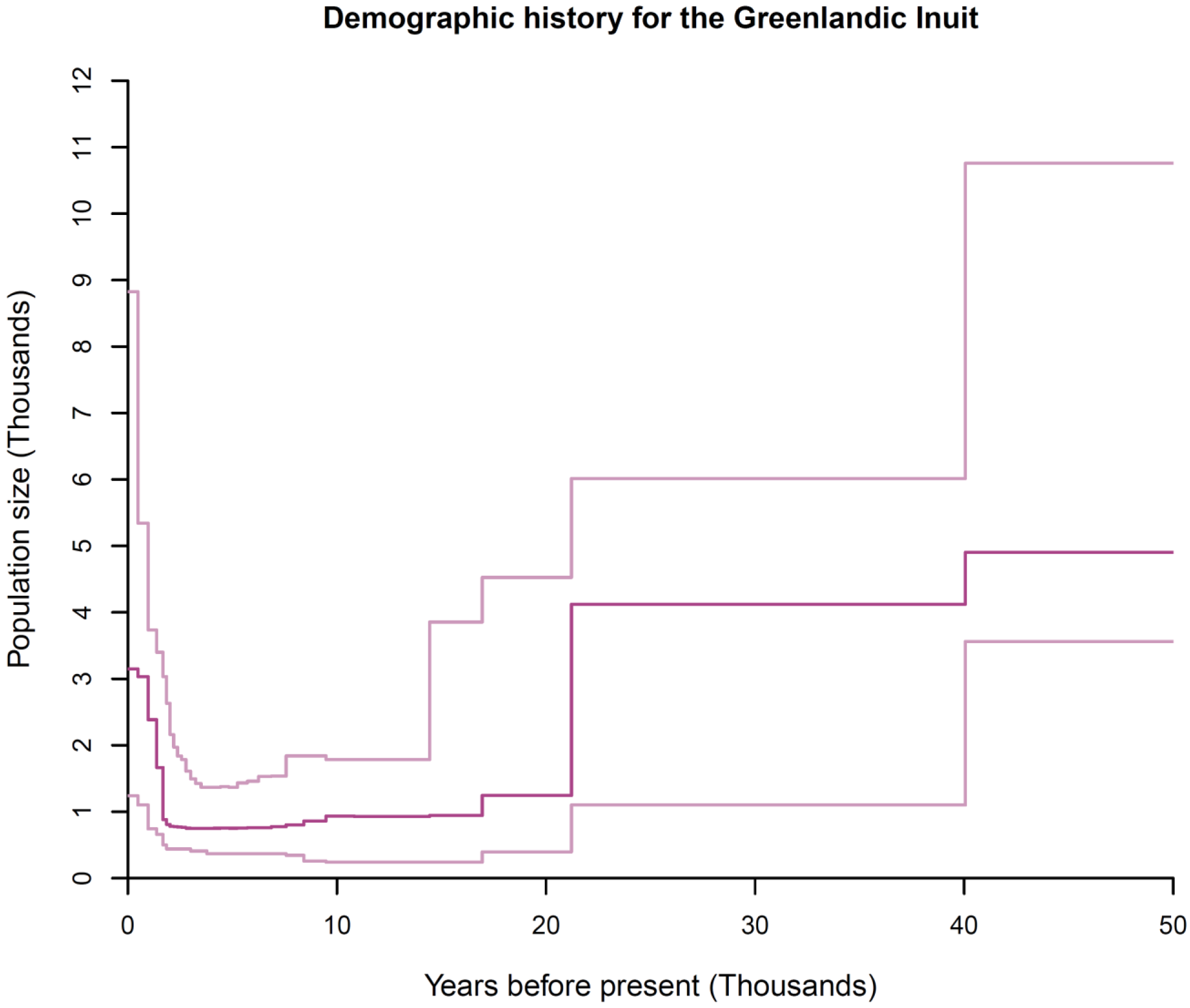
Stairway plot for the Greenlandic Inuit population. The dark pink line shows the estimated diploid population size changes in discrete increments for the last 50k years. The estimates were obtained with the method “Stairway plot”, which bases its estimates on the site frequency spectrum. The estimates are based on an assumption of a mutation rate of 1.2 × 10^−18^ per site per generation and a generation time of 24 years. The light pink lines represent 95% CI based on bootstraps. This analysis was based on 41.222.102 sites.

Next, we compared the Site Frequency Spectrum (SFS) of the GI population to the SFSs of a number of other human populations from the 1000 genomes project (Figure 2) and Cambodians and Mayans from Henn et al. ^5^ (Figure S8). Because the SFS is affected by demographic history ^28,29^, this comparison should reveal the extent to which the recent very long bottleneck has made the GI more extreme than other OOA populations. These comparisons show that the SFS for the GI population is flatter than for all the other examined populations, which is also reflected in markedly higher nucleotide diversity for the variable sites (π_var_), which is inversely correlated with π for all sites. Thus compared to other populations, the GI population has had both a larger depletion of rare variants, as well as a higher increase in allele frequencies for remaining variants. Hence the GI population shows evidence for having undergone a more extreme population bottleneck than the other populations, which means that analyses of this population are likely to be valuable in resolving the open questions about whether small effective population sizes have had an effect on genetic load and selection in human populations.

**Figure 2.**
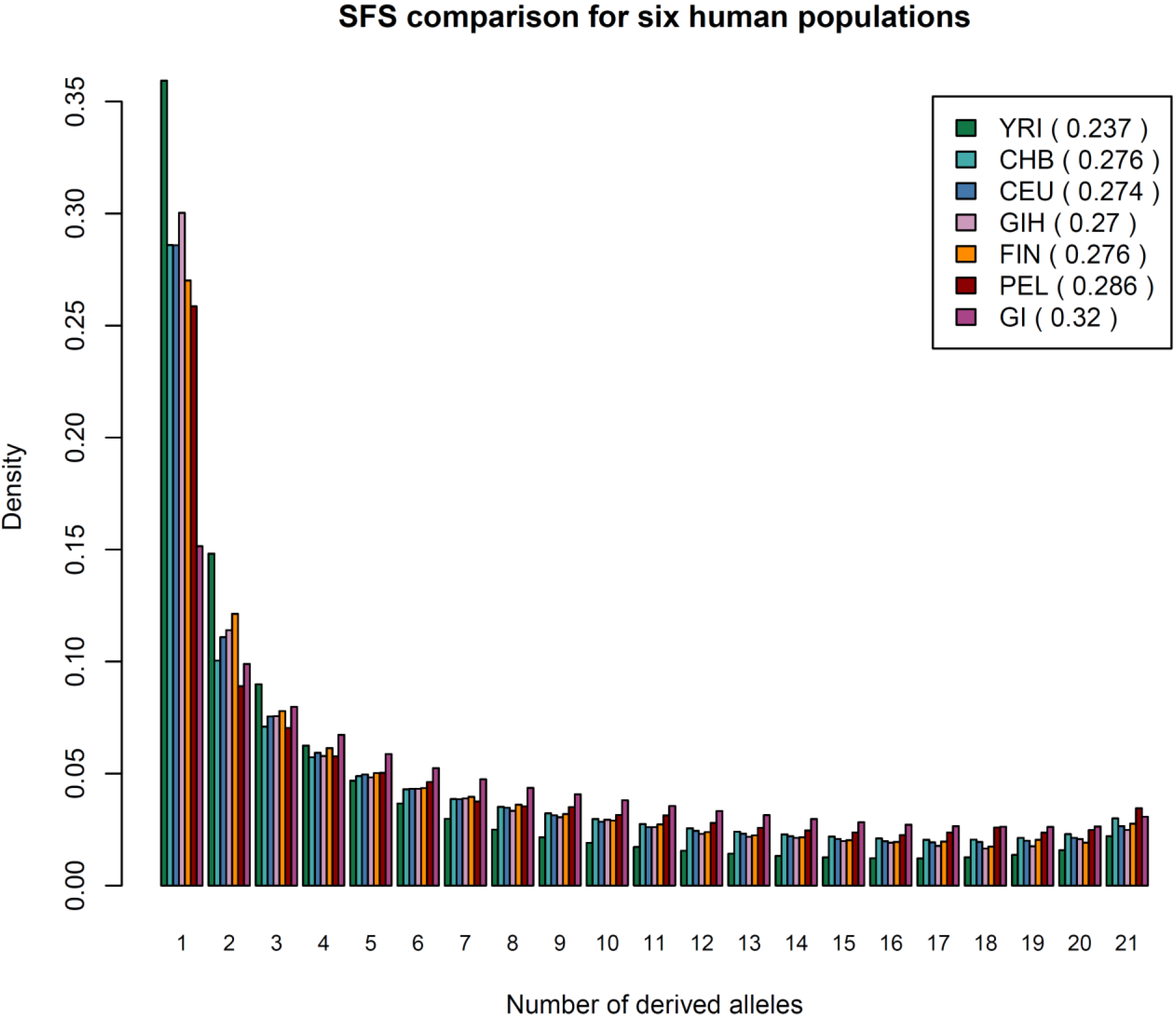
Site frequency spectrum for six human populations. We used 11 randomly sampled individuals from each population to infer the site frequency spectrum and excluded fixed categories. The GI population has fewer sites in the singleton category, but more in the remaining more “common” categories. Each population is followed by a π_var_ estimate per variable site.

### Comparison of genetic load in GI and CEU

Motivated by these initial findings, we compared the genetic load between GI and CEU to investigate to which extent, if any, the long history with a reduced population size has led to an increased genetic load in the GI population. For this comparison we used the exome data from the 18 GI combined with exome data from 18 individuals with European ancestry (CEU). Since there is no direct way to calculate load from genetic data, we examined several different statistics, most of which have recently been proposed as approximations to genetic load.

First, we examined load statistics that assume that all alleles have an additive effect. In particular, we looked at the average number of derived alleles per individual, which has previously been used both by Fu et al. ^9^ and Do et al. ^8^. We performed the comparison between GI and CEU by calculating the ratio of this statistic in GI and CEU. Here a ratio above one would indicate that the average number of derived alleles in GI is higher than that in CEU. We calculated this ratio both for all sites and for sites in 9 different subcategories that represent a range of different levels putative deleterious effect: five categories based on functional annotation and four based on GERP scores (for details see Material and Methods). We did not find a significant difference between the numbers of derived alleles per individual between the two populations for any of the different functional annotation-based categories, i.e. the ratios did not differ significantly from 1 (Table 1 and Figure 3, Figure S4, Figure S1, S2, Table S1 and Table S2). The ratio of derived alleles in the more deleterious GERP categories show a 1-4% increase in GI compared to CEU (Figure 4), but this difference is not significant. Similar results are obtained from the R_X/Y_ ratio from Do et al. ^8^. Hence, the results from both these statistics suggest that, under an additive effect model, the load is the same in the across populations (Table 1, Table S1 and Table S2). However, we note that the percentage of the derived alleles per individual that come from sites fixed for the derived allele differs markedly between the two populations. This is seen for all sites as well as in most functional subcategories (e.g. LoF alleles and GERP scores higher than 6; Table 1; column 4). We also note that the average number of derived alleles per SNV differs significantly between the two populations in all categories (Table 1; column 6). All the ratios are significantly larger than 1, indicating that per SNV, the derived allele is at higher frequency in GI compared to CEU. This observation combined with our previous observation that the average number of derived alleles per individual is the same in the two populations, reveals that the derived alleles in GI are from fewer sites with higher derived allele frequencies and thus that the load under an additive model is distributed differently in the two populations.

**Figure 3.**
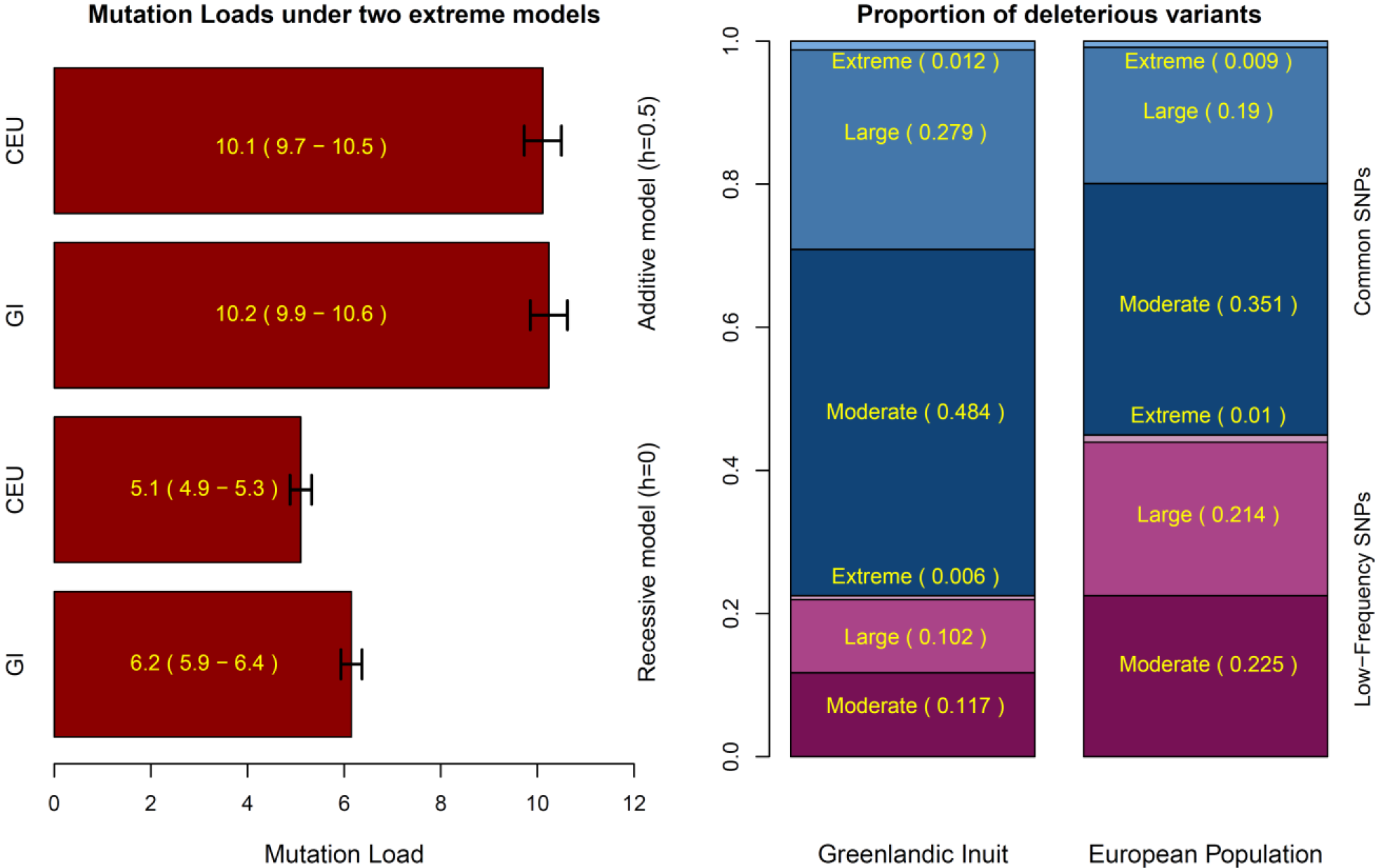
Mutation Load and Load proportions. The left plot shows the genetic load using a fully additive model (top two bars) and a fully recessive model (bottom two bars). These genetic load values are based on annotated GERP scores converted to selection coefficients using the approach from Boyko et al.^6^. We note that if the selection coefficients from Henn et al.^5^ are used instead, we see qualitatively similar results. The genetic load was calculated as in Lohmuller^11^. Black error bars indicate 95% C.I. The right plot shows the proportion of deleterious variants classified by GERP score (Moderate: 2<GERP<4, Large: 4<GERP<6, Extreme: 6<GERP) and their frequency in the populations (Low vs common). Low frequency SNPs are here defined as singletons and doubletons (equivalent of a frequency of at most 1/18≈0.056), while common SNPs are defined as tripletons or more than that including fixed derived sites (equivalent of a frequency of 1/12 or above).

**Figure 4.**
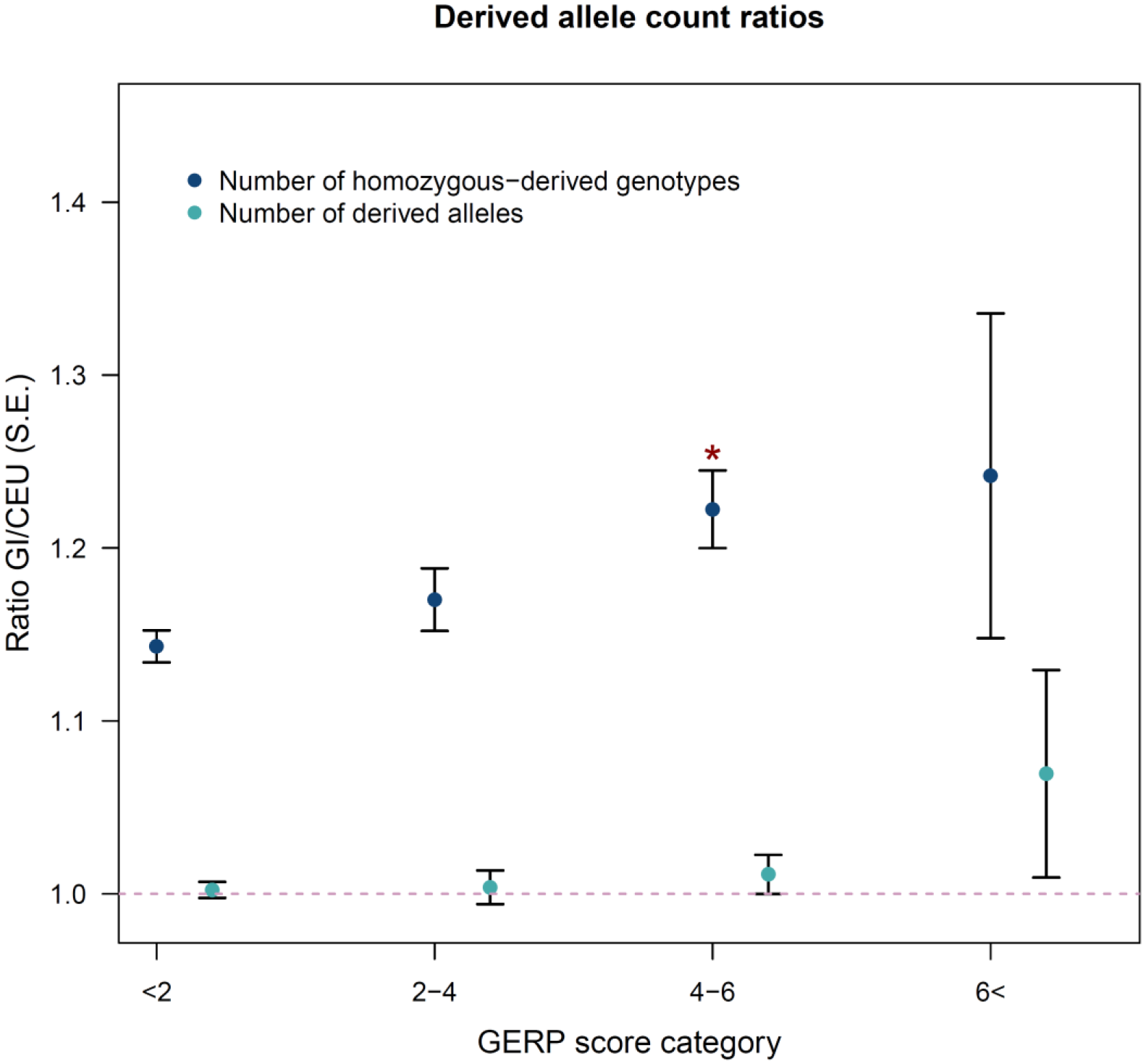
Load statistic ratios. For each of four GERP score categories two ratios calculated based on sites located in exons are shown: the ratio of derived allele counts in GI versus CEU (turquoise) and the ratio of homozygous-derived genotype counts in GI versus CEU (blue). The former can be viewed as an approximation for the ratio of load between the two populations under an additive model and latter can be viewed as an approximation for the ratio of load between the two populations under a recessive model. Standard error for each ratio is indicated by error bars. Additional Information is available in Table 1. The * indicates significance compared to the neutral GERP score category (GERP<2) (*P* = 5.3 × 10^-4^) using a z-test (compare column two for row 7 and 9 in Table 1).

**Table 1.** Summary of load investigation. Comparisons between 18 Greenlandic Inuit (GI) and 18 Utah Residents (CEPH) with Northern and Western Ancestry (CEU) individuals using 93.047 SNPs sites located in genic regions with full information for both populations. Ratios are the sum within GI divided by the sum within CEU. Standard errors are given in parentheses.

| Category (N SNV) | Ratio of derived allele counts (SE) | Ratio of homozygous-derived genotype counts (SE) | Ratio of derived alleles in fixed sites (SE) | Derived alleles from fixed sites | Ratio of derived alleles per SNV (SE) | R<sub>xy</sub> (SE) |
| --- | --- | --- | --- | --- | --- | --- |
| **Derived Alleles (49631)** | 1.002 (0.004) | 1.156 (0.008) | 4.413 (0.301) | GI:17.0% CEU:3.9% | 1.484 (0.009) | 1.004 (0.008) |
| **Synonymous (20693)** | 1.006 (0.005) | 1.156 (0.010) | 4.733 (0.420) | GI:18.2% CEU:3.9% | 1.420 (0.011) | 1.013 (0.012) |
| **NCE+UTR (4919)** | 1.006 (0.012) | 1.168 (0.022) | 5.268 (0.900) | GI:17.9% CEU:3.4% | 1.466 (0.023) | 1.013 (0.026) |
| **Missense (23582)** | 0.996 (0.006) | 1.151 (0.012) | 3.899 (0.377) | GI:15.5% CEU:4.0% | 1.550 (0.013) | 0.993 (0.013) |
| **LoF (400)** | 1.009 (0.063) | 1.296 (0.131) | 2.500 (2.840) | GI:9.3% CEU:3.8% | 1.625 (0.115) | 0.991 (0.104) |
| **LoF<sup>a</sup> (278)** | 1.045 (0.099) | 1.457 (0.286) | - | GI:0% CEU:4.4% | 2.026 (0.194) | 1.064 (0.142) |
| **GERP < 2 (29183)** | 0.999 (0.005) | 1.140 (0.009) | 4.293 (0.291) | GI:18.4% CEU:4.0% | 1.414 (0.009) | 0.999 (0.010) |
| **GERP >2:<4 (9853)** | 1.001 (0.010) | 1.169 (0.018) | 4.725 (0.757) | GI:15.3% CEU:3.1% | 1.517 (0.019) | 1.002 (0.019) |
| **GERP >4:<6 (10211)** | 1.011 (0.011) | 1.222 (0.022) | 4.659 (0.919) | GI:12.7% CEU:2.8% | 1.697 (0.023) | 1.021 (0.021) |
| **GERP >6 (353)** | 1.069 (0.060) | 1.244 (0.093) | - | GI:8.6% CEU:0% | 1.888 (0.130) | 1.127 (0.113) |

<sup>a</sup> = unambiguous sites (sites where all possible functional annotations were LOF annotations)
- indicates that too few sites fall in this the category to allow a meaningful result

Next, we aimed to compare the genetic load under the assumption that deleterious alleles have a recessive effect. To do this we examined the average number of homozygous derived genotypes per individual. The ratios of this statistic were significantly higher than 1 (Table 1; column 2 and Figure 3, Figure S4), indicating an accumulation of homozygous derived genotypes in the GI population. This was seen for all categories, even for the LoF categories. In fact, the ratio is significantly higher in sites with large deleterious effects, i.e. GERP scores above 4, compared to neutral sites, i.e. sites with GERP scores below 2 (see asterisk in Figure 4). These observations suggest that the load is increased in GI, if the mode of selection is recessive.

We also looked at the load statistic proposed by Henn et al. ^10^, which we denote as GERP score load. This statistic approximates load using the original definition by Kimura et al. ^1^, where each GERP score category is translated into a selection coefficient. We coupled these groups of selection coefficients with two models: one where *h* was set to 0. 5, i.e. assuming alleles have an additive effect and one where *h* was set to 0; i.e. assuming alleles have a recessive effect (for details see Materials and Methods). When doing so, we saw no significant difference between the two populations under the additive model (10.1 (S.E. 0.4) for CEU vs 10.2 (S.E. 0.4) for GI), but a 22% increase in the GERP score load under the completely recessive model (5.1 (S.E. 0.2) vs 6.2 (S.E. 0.3)) (Figure 3). Furthermore, putatively deleterious SNVs (GERP score > 2) that contribute to the genetic load are much more common in the GI population compared to the CEU population (Figure 3). Hence, the results obtained based on GERP scores lead to qualitatively the same conclusion as the other statistics.

We note that we also performed similar load comparisons to 18 individuals with East Asian ancestry (CHB) and the 18 individuals with African ancestry (YRI) to ensure that our conclusions were not artifacts of particular features of the CEU. In particular, we produced the equivalents of Table 1 and figure 4 for these two populations (Table S1, Table S2, Figure S1 and Figure S2). When doing so we reached the same conclusions for both populations with one exception: when comparing GI and CHB, the ratio of homozygous-derived genotype counts is indeed higher in sites with large deleterious effects, i.e. GERP scores above 4, compared to neutral sites, but not significantly so (Figure S1). However, for YRI this difference is significant (asterisk in Figure S2).

### Comparison of genetic load based on simulations

To investigate these results further, and try to assess what can be concluded from them, we performed simulations of GI and CHB populations using the demographic history of these two populations inferred in Fumagalli et al. ^16^. We performed the simulations of 12 different selection scenarios that varied both in the effect model used, i.e. *h*, and the selection coefficients, s, that reflect how deleterious the alleles are. Based on the simulated data we calculated the ratio of the number of derived alleles between the two populations and this ratio was not significantly different from 1 in any of the scenarios (Table S6).

We also counted the number of homozygous derived genotypes and, as expected, we observed a significantly higher number GI than in CHB (Figure S8-9). The increase is about 19% even without selection (Table S6).

Finally, we calculated the true load based on the true effect model and the true selection coefficient for each site. Interestingly the ratio of this true load in GI and CHB is significantly higher than 1 in all scenarios, indicating a higher load in GI than in CHB. This difference decreases as *h* increases, however, importantly, even in the scenarios where additive selection (*h*=0.5) was simulated, the ratio of loads were all significantly above 1, with the increase varying from 1 to 4% (Figure S9 and Table S6) depending on the strength of selection simulated. The highest load difference was observed for the scenario with *s*=0.002.

These results are interesting because they suggest that load statistics, like the number of derived alleles, will not necessarily reveal if there is a difference in load between two populations. Thus the results suggest that the fact that GI do not have higher additive load statistic than CEU in our data cannot necessarily be used to conclude that there is no difference in load even if all alleles have an additive effect. For the same reason, it cannot necessarily be used to conclude that selection has not been acting less efficiently in the GI.

### Effectiveness of selection

To investigate to what extent the long recent bottleneck in GI has affected the effectiveness of selection, we looked at two other summary statistics, which have both previously been used to address this question in the context of the OOA bottleneck: SFSs and the proportion of non-synonymous to synonymous mutations^13^.

We first compared SFSs for GI and the CEU. Specifically we made SFSs for each of five the functional categories also used in our previous analyses: synonymous, a combined category of non-coding exons (NCE) and the leading and trailing untranslated region (UTR), missense and finally two different LoF categories (Figure S3 and Figure S4). These SFSs show that GI have a larger proportion of alleles found in higher frequencies, not only overall, but also within all of the functional categories including LoF, and thus potentially highly deleterious alleles. And consistent with the results described previously, the SFSs also show that the GI population has a higher proportion of fixed putatively deleterious derived alleles in all the functional categories. In contrast, the CEU population has a clear skew towards rare variants. This difference, may reflect the effects of less efficient purifying selection in GI and/or recent growth in CEU ^6,29^. However, we note that based on simulations similar to the ones presented above, but where each deleterious allele is set to neutral (changing the selection coefficient to 0) in GI after the start of the GI bottleneck (Figure S10), we observed a fairly similar SFS to the one where selection continues to happen after the bottleneck. Thus based on the simulations we do not necessarily expect to be able to observe if selection is less effective based on large differences in the SFS of the different functional categories. Next, we compared the ratio of non-synonymous to synonymous SNPs in the GI populations to that of the CEU population using the Combined GI and CEU dataset, following what was done by Lohmueller et al. ^13^ (Table S4). Specifically, we tested for a difference in this ratio between the two populations both among all sites and among sites from different GERP scores categories. We found that ratio of the number of non-synonymous to synonymous SNPs is significantly higher in CEU than in GI, when considering all sites (first row in Table S4). When considering different GERP score categories, the non-synonymous to synonymous ratio in GI becomes even lower relative to that seen in CEU with increasing levels of deleteriousness.

We further investigated this pattern by defining neutral sites as those with GERP scores below 2 and deleterious variants as those with GERP scores above 4 and then calculating the ratio of deleterious to neutral SNPs for both GI and CEU. When doing so the deleterious to neutral SNP ratio is significantly lower in the GI compared to the CEU (Table S4; row 2) and is almost as low as the ratio of non-synonymous to synonymous sites among the most deleterious GERP scores (Table S4; row 5 and 6). Hence, there seem to be a higher proportion of deleterious variants in CEU than in GI, particularly for highly deleterious variants. This is compatible with the notion that selection has not been acting less efficiently in GI than in CEU. Hence the different statistics point towards different conclusions, which we will discuss later.

### Consequences for disease mapping

Regardless of whether selection has acted less efficiently in GI or not, our results clearly show that individual deleterious variants tend to have higher frequency in GI compared to CEU (Figure 3). This is potentially of high importance for quantitative trait mapping and disease mapping as it means that mapping in the GI population will in some cases be more powerful in the GI population than the CEU population ^30^ just like it is well-known to be the case for other historically isolated populations like the Finnish population and the Native Americans. To investigate the potential of GI, and compare it to that of other populations, we divided alleles into bins according to their frequency in larger reference populations and for each bin determined how often the alleles are common among GI compared to how often they common in other populations. To do this we used allele frequencies from the Exome aggregation consortium (ExAC). As can be seen in Figure 5 and Figure S9-S11, we find that rare alleles among Non-Finnish Europeans are more likely to be common in GI than in Finland, East Asia and Latinos. This pattern is especially pronounced for variants that are extremely rare in Non-Finnish Europeans (MAF < 1×10^−3^). Similar patterns are seen when looking at variants that are rare in East Asians. This suggests that the GI population does indeed have potential to provide increased power to detect alleles that are rare in Non-Finnish Europeans and East Asians. And it suggests that this potential is even bigger than that of both Latinos and Finns; populations that are considered particularly powerful for mapping.

**Figure 5.**
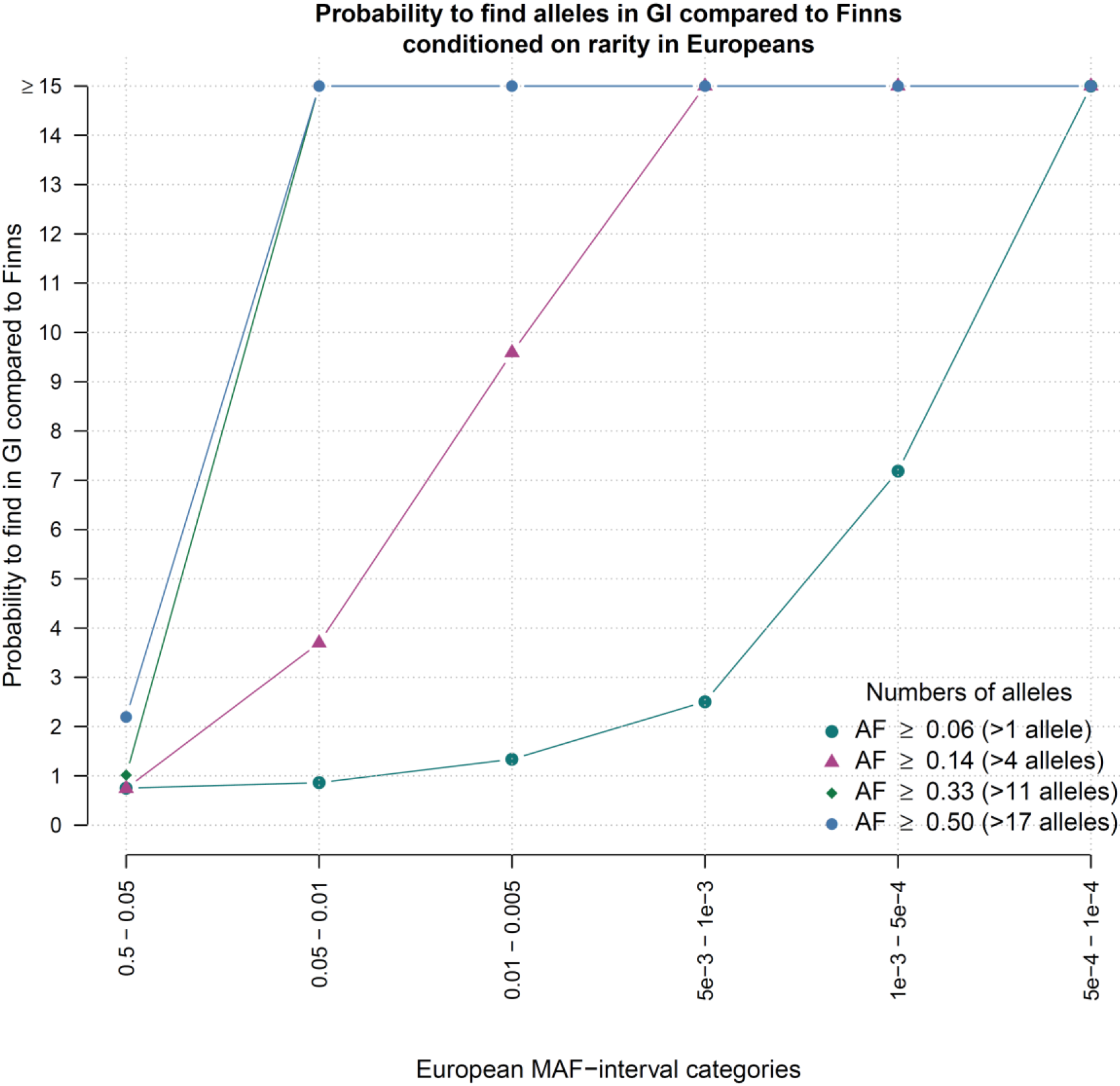
Probability of common alleles. For alleles in different frequency categories in Europeans (x-axis) the points show how much more likely the alleles are to be common in GI than in Finns (y-axis). Common is here defined in 4 different ways, each represented by a specific color point, e.g. the dark blue circles represents results for analyses made with common defined as more than 2 out of 36 alleles, which corresponds to a frequency of ≈0.056 (for the remaining definitions, see the figure legend).

To pursue this potential of the GI population in a simple manner we investigated all the LoF including frameshift variants that are present in at least 2 copies in the GI dataset and are rare, but not absent, in both the European and EAS populations (MAF below 0.5%) (Table 2, Table S5). We did this because such alleles are of potential interest for the Europeans and EAS, but difficult to map and investigate in these populations. This approach led to the identification of 6 SNVs and 14 indels. One of the SNVs, located in *TBC1D4*, was recently shown to have a very large impact on type 2 diabetes^15^ and one of the indels, located in *SI*, was recently shown to have large impact on a sucrase-isomaltase deficiency ^31^. Other interesting variants include a SNV in *SEMA4C* which gene is associated with neonatal lethality^17,32^ and SNVs in *CRYGA* and *USP45* which might be involved in cataract and DNA repair, respectively ^33,34^.

**Table 2.** Summary of Loss of Function mutation. LoF variant (Stop & splice variants) annotation for alleles that are rare in Europeans and East Asians (below a MAF of 0.5%) but not lost.

| rs number | Gene name | Possible clinical implications | Genotypes among the 18 Inuit (Parents only) | Reference allele |
| --- | --- | --- | --- | --- |
| rs12471298 | *SEMA4C* | Neonatal lethality | CC=12,CA=6 | C |
| rs61736969 | *TBC1D4* | Type 2 diabetes | GG=7,GA=7,AA=4 | G |
| rs116344874 | *CRYGA* | Cataract | GG=13,GA=5 | G |
| rs189281869 | *USP45* | DNA repair (following UV irradiation) | TT=15,TA=3 | T |
| rs189664399 | *SIGLEC7* | unknown | CC=15,CT=3 | C |
| rs201985967 | *SPAG4* | unknown | TT=13,TA=5 | T |

## Discussion

We have performed analyses of exome data from the GI population with the aim of investigating what consequences small population size has had on genetic variation in humans. We first investigated the demographic history of the GI population and found that it dramatically decreased in size ~23.000 years ago and remained small for more than 20 thousand years, which corroborates previously reported results ^16^. Furthermore, we observed a flattening of the SFS for populations with increasing distance to Africa, and that the SFS for the GI population was even more extreme than those for the equally distant Mayan and Peruvian populations. This suggests that the GI population underwent a strong bottleneck after it split from the Native Americans. This is consistent with our estimates of the GI population size over time (Figure 1), where the bottleneck persisted long after the split from Native Americans. This finding makes the GI population highly relevant to the study of how demography has shaped deleterious variation in humans.

Motivated by this we investigated whether the severe bottleneck has affected the genetic load of the GI population by comparing it to CEU. In this context, it is important to re-iterate that genetic load cannot be directly calculated from exome data ^11^. It can only be approximated through predictions of whether and to what extent alleles are deleterious, e.g. via functional annotation. Also, the approximations depend on the underlying effect model, i.e. whether the effect of the deleterious alleles is additive or recessive. We therefore compared the genetic load between the GI and CEU populations using several different proxy statistics. Our analyses reveal no significant differences between the two populations when using proxy statistics that assume an additive effect, like the number of derived alleles, but they do show large significant differences when using proxy statistics for load that assume a recessive effect, like the number of homozygous-derived genotypes (Table 1). Importantly, we observed similar results when comparing GI to CHB and YRI (Table S1 and Table S2). These observations are in line with the results from several previous studies ^7,8^ and are consistent with our simulations where we observe no significant effect on the number of derived alleles and a large significant difference in number of homozygous derived genotypes (Figure 4, Figure S1, Figure S2, Figure S9 and Table S6). When looking at sites with GERP scores or functional annotations, which suggest that that they harbor highly deleterious alleles we observed a larger increase in number of derived alleles in GI than in CEU, although the increase is small and nonsignificant using jackknife resampling. Therefore, our results are not necessarily conflicting with Henn et al. ^5^ that found that the number of deleterious alleles increases with distance from Africa. However, we would have expected to see a stronger difference in GI than in the Mayan used by Henn et al. ^5^. Though, it is possible that we would see such evidence in our ratios if we analyzed a larger number of samples.

Interestingly, our simulations show a significant increase in the true load in the simulated GI individuals compared to the simulated CHB individuals. Even under an additive model, there is a small but significant increase of 1-4% depending on the selection coefficient. This suggests that small population size over a long period of time can indeed lead to increased load even if all alleles have an additive effect. These simulation results are in conflict with simulation-based conclusions from Simons et al. ^7^, based on deleterious alleles counts. However, our results fit very well with recent results from Harris and Nielsen^35^: when simulating Neanderthals and humans and assuming a severe bottleneck in Neanderthal that lasted many times longer than the bottleneck in GI, they observed at large significant decrease in fitness among Neanderthals even under an additive effect model. Importantly, our simulation results also revealed that the significant increase in load observed under additive selection, did not lead to a significant difference in the number of derived alleles. This suggests that the fact that we do not see a significant increase in the number of derived alleles in GI compared to CEU (or the other two populations) does not necessarily mean that there is no difference in load under an additive model. Furthermore, it supports the conclusion from Lohmueller ^11^ that part of the reason why different studies reach different conclusions about load is that they are based on different approximations of load – some of which may be far from ideal.

Either way, our results combined with previous results suggest, that if there are increases in genetic load under an additive model due to the small population size and bottlenecks, these increases are small even for an extreme population like the GI. Our results also clearly show that the extent to which the load is increased in GI due to the severe bottleneck depends heavily on the true underlying effect model. Hence to fully answer this question more knowledge about the true effect model is needed.

The above conclusions make it difficult to use comparisons of our overall load estimates to make conclusions about whether or not the severe bottleneck in GI has led to less efficient selection. We therefore also looked at other statistics to address that question. One of these was the ratio of non-synonymous to synonymous SNPs. If selection has acted less effectively in GI compared to larger populations like CEU then we would expect this ratio to be higher in the GI population than in CEU. We observe the opposite, which could suggest that selection has not acted less efficiently in GI. However, we note that the ratio of non-synonymous to synonymous SNPs is highly sensitive to demographic changes and that our observations could also be explained by the recent explosive population growth in CEU ^11^. Importantly, we also note that if selection has acted equally efficiently in the two populations, then we would expect that the ratio of homozygous-derived genotypes in the two populations would be the same across categories with different levels of deleterious effect. However, we find that this is not the case. On the contrary, the ratio of homozygous-derived genotypes is significantly higher in sites with large deleterious effects, i.e. GERP scores above 4 and below 6 as compared to neutral sites, i.e. sites with GERP scores below 2 (see asterisk in Figure 4). This result was also seen when comparing GI and YRI (asterisk in Figure S2) but not when comparing GI and CHB (see Figure S1). In the latter case there is an increase in the ratio between the two categories but it is not significant (Figure S1). This suggests that the impact of selection is not the same in GI compared to the YRI and CEU populations and potentially not CHB either. More specifically, it suggests that selection against deleterious alleles has acted less efficiently in GI under a recessive model compared to the other populations. This argument is supported by the fact that we observed a similar pattern in our simulations. Here the increase in the homozygous-derived genotype ratio for the sites under selection (denoted as load ratio in Supplementary Table 6) under a recessive model is particularly large for alleles with selection coefficient *s*=0.002, which correspond to a GERP score of 4-6 ^5^(Figure S9-S11). This load difference would also explain the higher allele frequencies of deleterious alleles frequencies in GI compared to in CEU (Figure 3). Hence these results combined suggest that selection may indeed have acted less efficiently, and that the higher ratio of synonymous to non-synonymous sites GI is most likely explained by the population growth in CEU as has been suggested in other studies as well^12^.

Load is a somewhat abstract concept, none-the-less it has very concrete connections to disease and although our comparisons of load in GI and CEU may not be entirely conclusive, they have clear consequences in the context of disease. Our load comparisons revealed that a lot of variants are lost, also deleterious variants, and that the deleterious alleles that are not lost are in higher frequency in the GI population than in the CEU population. This has several disease related implications for the Greenlandic population. First, it has implication for disease risk. For complex disorders, where many loci are involved, we do not expect a large difference in genetic load and thus disease risk. This can be illustrated by the *TBC1D4* variant’s impact on the type 2 diabetes prevalence. Despite its large effect and relatively high frequency ^17^ (23% MAF in GI) the Greenlandic population has not had a historically high incidence of type 2 diabetes in Greenland ^36^. We suspect that the absence of other type 2 diabetes variants in GI are compensating for the presence of the common variants, but acknowledge that other variants may be found if we investigated a larger amount of data. However, for the rarer more monogenetic traits, we suspect that the genetic load, and thus disease risk, will be very different in Greenland. Because fewer variants are involved, the variance of the prevalence of the trait will be much greater. This means that such diseases will likely either entirely absent from the population or be more prevalent. The SI frameshift variations effect on sucrase-isomaltase deficiency is an example of the latter. SI deficiency is a very rare disorder in large populations, but is estimated to affect 5-10% of individuals in Greenland ^37,38^ This is presumably solely due to a single SI frameshift variant, which we find in a homozygous state in 3 of the individuals among the 9 GI trios (Table S5).

It also has implication for our ability to characterize the function of certain genes, including disease related genes. Due to the higher frequency of deleterious alleles, even LoF alleles, the GI are enriched for homozygous functional knockouts compared to the CEU. This enables investigation of the function of the genes that harbor such mutations.

Finally, it has implication for disease mapping. It has long been acknowledged that populations, which have undergone a recent population bottleneck and therefore, like the Greenlandic population, carry deleterious alleles with higher allele frequency compared to larger populations are useful in disease studies, because the increased allele frequency (along with increased LD) leads to increased power in association testing. The Finnish population is one such population, where two novel mutations have been found to be associated with lipoprotein levels ^39^. Other examples included various Native American populations where studies have also led to the detection of several novel disease variants. Interestingly, our analyses show that GI outperform both Finns and Latinos in terms of the chance of providing improved power in association testing due to higher allele frequencies (Figure 5, Figure S11-S14). Thus likely, studies aiming to find novel association will have even better power to do so in the GI population compared to previously studied isolated populations.

Our simple screening of the LoF variants provides a simple demonstration how useful GI can be for disease mapping. It also clearly shows that studies of the Greenlandic population can be used to identify alleles of large effect. However, populations like the Greenlandic are also useful for identifying variants with lower effect sizes, because, as this study shows, such alleles are much more likely to be of high frequency than are alleles of large effect. Importantly, since our analyses of allele frequencies among GI compared to Finns and Latinos were carried out for alleles that are rare, but indeed present among Europeans and East Asians our results also suggests that disease mapping in the Greenlandic population has great potential to lead to identification of variants that are also important in larger populations, like the Europeans and East Asians. Thus, all in all our results show that studies of GI population constitute a particularly promising approach in future disease mapping.

## Supplemental Material

Our supplemental data comprise 14 figures and 6 tables.

## Acknowledgements

The Novo Nordisk Foundation Center for Basic Metabolic Research is an independent Research Center at the University of Copenhagen partially funded by an unrestricted donation from the Novo Nordisk Foundation (http.metabol.ku.dk).

## Web Resources

Henn et al.^5^ https://ecoevo.stonybrook.edu/hennlab/data-software/

ExAC browser ftp://ftp.broadinstitute.org/pub/ExACrelease/release0.3/ExAC.r0.3.sites.vep.vcf.gz

GERP scores http://hgdownload.cse.ucsc.edu/gbdb/hg19/bbi/Allhg19RS.bw

## References

1. Kimura, M., Maruyama, T., and Crow, J.F. (1963). Mutation load in small populations. Genetics 48, 1303–1312.

2. Ohta, T. (1973). Slightly deleterious mutant substitutions in evolution. Nature 246, 96–98.

3. Charlesworth, B. (2009). Fundamental concepts in genetics: effective population size and patterns of molecular evolution and variation. Nat. Rev. Genet. 10, 195–205.

4. Morton, N.E., Crow, J.F., and Muller, H.J. (1956). An estimate of the mutational damage in man from data on consanguineous marriages. Proc Natl Acad Sci U S A 42, 855–863.

5. Henn, B.M., Botigué, L.R., Peischl, S., Dupanloup, I., Lipatov, M., Maples, B.K., Martin, A.R., Musharoff, S., Cann, H., Snyder, M.P., et al. (2015). Distance from sub-Saharan Africa predicts mutational load in diverse human genomes. Proc. Natl. Acad. Sci. 113, E440–E444.

6. Boyko, A.R., Williamson, S.H., Indap, A.R., Degenhardt, J.D., Hernandez, R.D., Lohmueller, K.E., Adams, M.D., Schmidt, S., Sninsky, J.J., Sunyaev, S.R., et al. (2008). Assessing the evolutionary impact of amino acid mutations in the human genome. PLoS Genet. 4, e1000083.

7. Simons, Y.B., Turchin, M.C., Pritchard, J.K., and Sella, G. (2014). The deleterious mutation load is insensitive to recent population history. Nat. Genet. 46, 220–224.

8. Do, R., Balick, D., Li, H., Adzhubei, I., Sunyaev, S., and Reich, D. (2015). No evidence that selection has been less effective at removing deleterious mutations in Europeans than in Africans. Nat. Genet. 47, 126–131.

9. Fu, W., Gittelman, R.M., Bamshad, M.J., and Akey, J.M. (2014). Characteristics of neutral and deleterious protein-coding variation among individuals and populations. Am. J. Hum. Genet. 95, 421–436.

10. Henn, B.M., Botigué, L.R., Bustamante, C.D., Clark, A.G., and Gravel, S. (2015). Estimating the mutation load in human genomes. Nat. Rev. Genet. 16, 333–343.

11. Lohmueller, K.E. (2014). The distribution of deleterious genetic variation in human populations. Curr. Opin. Genet. Dev. 29, 139–146.

12. Lohmueller, K.E. (2014). The Impact of population demography and selection on the genetic architecture of complex traits. PLoS Genet. 10, e1004379.

13. Lohmueller, K.E., Indap, A.R., Schmidt, S., Boyko, A.R., Hernandez, R.D., Hubisz, M.J., Sninsky, J.J., White, T.J., Sunyaev, S.R., Nielsen, R., et al. (2008). Proportionally more deleterious genetic variation in European than in African populations. Nature 451, 994–997.

14. Raghavan, M., DeGiorgio, M., Albrechtsen, A., Moltke, I., Skoglund, P., Korneliussen, T.S., Gronnøw, B., Appelt, M., Gulløv, H.C., Friesen, M., et al. (2014). The genetic prehistory of the new world arctic. Science 345, 1255832.

15. Gulløv, H.C. Grønlands forhistorie (Gyldendal, Copenhagen, 2004).

16. Fumagalli, M., Moltke, I., Grarup, N., Racimo, F., Bjerregaard, P., Jørgensen, M.E., Korneliussen, T.S., Gerbault, P., Skotte, L., Linneberg, A., et al. (2015). Greenlandic Inuit show genetic signatures of diet and climate adaptation. Science 349, 1343–1347.

17. Moltke, I., Grarup, N., Jørgensen, M.E., Bjerregaard, P., Treebak, J.T., Fumagalli, M., Korneliussen, T.S., Andersen, M.A., Nielsen, T.S., Krarup, N.T., et al. (2014). A common Greenlandic TBC1D4 variant confers muscle insulin resistance and type 2 diabetes. Nature 512, 190–193.

18. McVean, G.A., Altshuler (Co-Chair), D.M., Durbin (Co-Chair), R.M., Abecasis, G.R., Bentley, D.R., Chakravarti, A., Clark, A.G., Donnelly, P., Eichler, E.E., Flicek, P., et al. (2012). An integrated map of genetic variation from 1,092 human genomes. Nature 491, 56–65.

19. Moltke, I., Fumagalli, M., Korneliussen, T.S., Crawford, J.E., Bjerregaard, P., Jørgensen, M.E., Grarup, N., Gulløv, H.C., Linneberg, A., Pedersen, O., et al. (2015). Uncovering the genetic history of the present-day greenlandic population. Am. J. Hum. Genet. 96, 54–69.

20. Li, H., Handsaker, B., Wysoker, A., Fennell, T., Ruan, J., Homer, N., Marth, G., Abecasis, G., Durbin, R., and Subgroup, 1000 Genome Project Data Processing (2009). The sequence alignment/map format and SAMtools. Bioinformatics 25, 2078–2079.

21. Danecek, P., Auton, A., Abecasis, G., Albers, C.A., Banks, E., DePristo, M.A., Handsaker, R.E., Lunter, G., Marth, G.T., Sherry, S.T., et al. (2011). The variant call format and VCFtools. Bioinformatics 27, 2156–2158.

22. Korneliussen, T., Albrechtsen, A., and Nielsen, R. (2014). ANGSD: Analysis of Next Generation Sequencing Data. BMC Bioinformatics 15, 356.

23. Cooper, G.M., Stone, E.A., Asimenos, G., Green, E.D., Batzoglou, S., and Sidow, A. (2005). Distribution and intensity of constraint in mammalian genomic sequence. Genome Res. 15, 901–913.

24. Goode, D.L., Cooper, G.M., Schmutz, J., Dickson, M., Gonzales, E., Tsai, M., Karra, K., Davydov, E., Batzoglou, S., Myers, R.M., et al. (2010). Evolutionary constraint facilitates interpretation of genetic variation in resequenced human genomes. Genome Res. 20, 301–310.

25. Liu, X., and Fu, Y.-X. (2015). Exploring population size changes using SNP frequency spectra. Nat. Genet. 47, 555–559.

26. Busing, F.M.T.A., Meijer, E., and Leeden, R.V.D. (1999). Delete-m Jackknife for Unequal m. Stat. Comput. 9, 3–8.

27. Messer, P.W. (2013). SLiM: Simulating Evolution with Selection and Linkage. Genetics 194, 1037–1039.

28. Marth, G.T., Czabarka, E., Murvai, J., and Sherry, S.T. (2004). The Allele Frequency Spectrum in Genome-Wide Human Variation Data Reveals Signals of Differential Demographic History in Three Large World Populations. Genetics 166, 351–372.

29. Nielsen, R., Bustamante, C., Clark, A.G., Glanowski, S., Sackton, T.B., Hubisz, M.J., Fledel-Alon, A., Tanenbaum, D.M., Civello, D., White, T.J., et al. (2005). A scan for positively selected genes in the genomes of humans and chimpanzees. PLoS Biol. 3, e170.

30. Hong, E.P., and Park, J.W. (2012). Sample Size and Statistical Power Calculation in Genetic Association Studies. Genomics Inform. 10, 117.

31. Marcadier, J.L., Boland, M., Scott, C.R., Issa, K., Wu, Z., Mcintyre, A.D., Hegele, R.A., Geraghty, M.T., and Lines, M.A. (2014). Congenital sucrase—isomaltase deficiency: identification of a common Inuit founder mutation. Can. Med. Assoc. J. 187, 102–107.

32. Maier, V., Jolicoeur, C., Rayburn, H., Takegahara, N., Kumanogoh, A., Kikutani, H., Tessier-Lavigne, M., Wurst, W., and Friedel, R.H. (2011). Semaphorin 4C and 4G are ligands of Plexin-B2 required in cerebellar development. Mol. Cell. Neurosci. 46, 419–431.

33. Kapur, S., Mehra, S., Gajjar, D., Vasavada, A., Kapoor, M., Sharad, S., Alapure, B., and Rajkumar, S. (2009). Analysis of single nucleotide polymorphisms of CRyGa and CRYGB genes in control population of western Indian origin. Indian J. Ophthalmol. 57, 197–201.

34. Perez-Oliva, A.B., Lachaud, C., Szyniarowski, P., Muñoz, I., Macartney, T., Hickson, I., Rouse, J., and Alessi, D.R. (2015). USP 45 deubiquitylase controls ERCC 1 - XPF endonuclease-mediated DNA damage responses. eMbO J. 34, 326–343.

35. Harris, K., and Nielsen, R. (2016). The genetic cost of neanderthal introgression. Genetics Early online April 2, 2016; DOI: 10.1534/genetics.

36. Jorgensen, M.E., Bjeregaard, P., Borch-Johnsen, K., Backer, V., Becker, U., Jorgensen, T., and Mulvad, G. (2002). Diabetes and impaired glucose tolerance among the Inuit population of greenland. Diabetes Care 25, 1766–1771.

37. McNair, A., Gudmand-Hoyer, E., Jarnum, S., and Orrild, L. (1972). Sucrose malabsorption in Greenland. Br. Med. J. 2, 19–21.

38. Gudmand-Høyer, E., Fenger, H.J., Kern-Hansen, P., and Madsen, P.R. (1987). Sucrase deficiency in Greenland. Incidence and genetic aspects. Scand. J. Gastroenterol. 22, 24–28.

39. Lim, E.T., Würtz, P., Havulinna, A.S., Palta, P., Tukiainen, T., Rehnström, K., Esko, T., Magi, R., Inouye, M., Lappalainen, T., et al. (2014). Distribution and medical impact of loss-of-function variants in the Finnish founder population. PLoS Genet. 10, e1004494.

